# Hematological parameters of high and low altitude Tibetan populations

**DOI:** 10.1101/2020.11.15.383752

**Authors:** Nipa Basak, Tsering Norboo, Mohammed S. Mustak, Kumarasamy Thangaraj

## Abstract

High altitude hypoxia is believed to be experienced at elevations more than 2500 meters. A few studies have shed light on the biochemical aspects of high altitude acclimatization that profoundly included the subjects sojourning to the high altitude from low altitude and observation of the transient changes. However, information regarding the difference between the adapted people in high altitude and their counterpart, who reside in the low altitude are lacking. To address that issue, we have measured various hematological parameters and level of serum erythropoietin (EPO) in Tibetan population, who are residing in both high and low altitudes. We observed significant difference (p value < 0.0001) between high and low altitude Tibetan, in various hematological parameters, including red blood cells (RBC) count, hematocrit (HCT) or packed cell volume (PCV), and hemoglobin concentration (Hb). In case of mean corpuscular volume (MCV), significant difference was observed only in females (p value < 0.0001). Mean corpuscular hemoglobin concentration (MCHC) was significantly different between both males and females, but age was a potential confounder. There was no significant difference in serum EPO level between these two groups, either in males or females, which might be due to blunted erythropoietin response in the Tibetan population. We have also analyzed correlation between serum EPO with Hb and serum EPO with HCT and found no significant correlation. In multiple regression analysis, low altitude and male-gender showed significant impact on both Hb and HCT. In conclusion, our study suggests significant perturbation of hematological parameters, when native high altitude populations migrated to low altitude and inhabited for a long period.

## Introduction

High altitude is clinically defined at altitudes ≥2500 meters above sea level, where physiological changes start appearing in vulnerable people ^1^. Extremely different environment in the high altitude area leads to ample changes in the population. A few studies have looked into the genetic aspects of high altitude adaptation ^2–5^ and physiological changes, however, physiological aspects of high altitude acclimatization studies performed till date profoundly included subjects travelling to the high altitude from low altitude; representing transient changes primarily or intermittent exposure to high altitude ^6–8^ or comparison of high altitude natives with non-native residents of high-altitudes ^9,10^ Adaptation is slow process and might take thousands of years to integrate genetic changes, whereas acclimatization can happen in hours/days to combat the stress in urgent manner. Hence, to observe the impact of environmental changes for few years, studying various physiological parameters or epigenetic parameters could be of better interest. Especially, differences in the people belonging to the same ethnic group, but inhabited in high and low altitudes could be interesting as these kinds of studies are rare and can give clarification of the environmental shift. In this study, hence, we have included Tibetan people who are native to high altitude since generations and Tibetan people, who are residing in low altitude for 50-60 years. Our high altitude group comes from different villages of Ladakh (altitude ≥ 4500 meters), India, whereas, low altitude group comes from Karnataka state, India (altitude ~ 1050 meters). They have migrated to Karnataka about 60 years ago, during Tibetan exodus event at the end of 1950’s from Tibet, when People’s Liberation Army (PLA) attacked Lhasa. They mainly migrated *via* Nepal and settled in Himachal Pradesh and Karnataka ^11,12^. Change of hematological parameters has been shown to be associated with various pathological conditions like myocardial infarction, leukemia, coronary heart disease, cardiovascular disease etc. ^13–16^ in earlier studies. In this study we aimed to investigate, how the hematological parameters, which includes white blood cells (WBC), red blood cells (RBC) count, hematocrit (HCT), packed cell volume (PCV), hemoglobin concentration (Hb), mean corpuscular volume (MCV), mean corpuscular hemoglobin (MCH) mean corpuscular hemoglobin concentration (MCHC), serum erythropoietin (EPO) level change in case of extreme environmental change i.e altitude in the population of same ethnic background.

## Materials and methods

### Subjects and study design

Our study includes a total of 79 blood samples of the high altitude Tibetans, who are residing at the altitude of ≥ 4500 meters above the sea level from different villages of Ladakh, India; and 89 individuals are low altitude Tibetans, who have migrated from Tibet to the southern state of Karnataka, India ^11,12^ about 50-60 years ago (Figure 1). Informed written consent was taken from all the participants and the Institutional Ethical Committee (IEC) of CSIR-Centre for Cellular and Molecular Biology, Hyderabad; and Ladakh Institute of Prevention; Ladakh approved this study.

**Figure 1:**
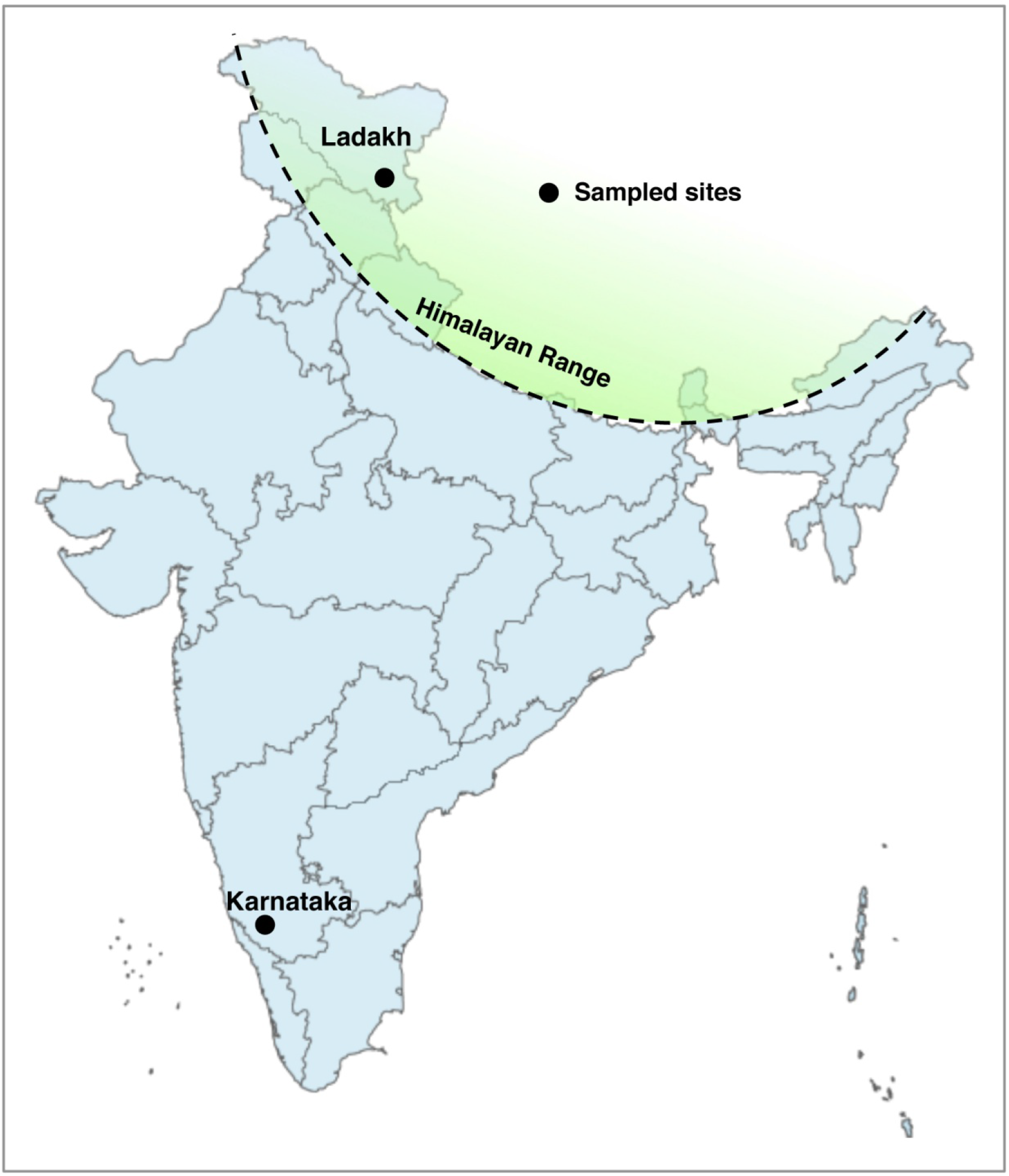
Sample collection sites, shown in the Indian map

About 3 ml of venous blood was collected in BD Vacutainer^®^ EDTA Tubes for hematological parameters assessment. About 3 ml of blood was also collected in BD Vacutainer^®^ Serum Tube. It was kept in rest at room temperature for half an hour. Clot was removed by centrifuging at 2,000 x g for 10 minutes. Clear serum was immediately transferred into clean polypropylene tubes and stored in dry ice temporarily and finally in −80 °C for further experiments.

### Anthropometric and hematological data

Age, gender, height, weight data were noted from all the subjects. Hematological data were acquired using automated hematology analyzer (ADVIA 2120i, Siemens AG, Germany) for the low altitude samples. For high altitude samples, hematological data were obtained using manual techniques. We considered WBC, RBC, HCT, MCV, Hb, MCHC, parameters for our study.

We could not measure the CBC profiles of the high-altitude and low altitude individuals, in the same method because of geographical differences of the places of sample collection. CBC profiles for the high altitude samples were obtained using manual techniques; while for the low altitude samples, these were obtained using automated hematology analyzer. However, both of the methods are known to be correlated well ^17,18^.

Quantikine human erythropoietin kit from R & D Systems (Minneapolis, MN, USA) was used to check serum EPO level in high and low altitude Tibetans.

### Data analysis

Graphpad Prism version 8.4.3 (San Diego, CA) was utilized for statistical analysis. D’Agostino-Pearson omnibus test was performed to check the normality of distribution of the dependent variables. Unpaired t-test and Mann-Whitney tests were performed for comparison between two groups, residuals of which followed Gaussian distribution and did not follow Gaussian distribution, respectively. For comparing more than two groups, One-way ANOVA and Kruskal-Wallis tests were performed. For multiple comparisons, Tukey’s and Dunn’s post hoc tests were performed. Spearman correlation coefficients were calculated to assess the correlation between parameters. Multiple linear regression analysis was performed in R package (version 3.6.1).

## Results

### Anthropometric and hematological parameters

Total participants in our study were 168 with age ranging from 17 to 92 years. The median age was 58 years. The BMI of the participants was in the range of 14.24 - 34.48 having median BMI 23.82. Table 1 shows, participants from high-altitude were having lower mean age and BMI, compared to low altitude participants.

**Table 1:** Description of anthropometric parameters of the subjects (n=168). Data has been expressed in mean ± SD format.

| Parameter | High altitude | Low altitude | Test, p value |
| --- | --- | --- | --- |
| Age (Years) | 49.32 ± 13.70 | 67.15 ± 18.42 | Mann Whitney test,<br><0.0001 |
| BMI (kg/m<sup>2</sup>) | 22.75 ± 3.67 | 25.28 ± 4.78 | Unpaired t-test,<br>0.0003 |

Since age and BMI was significantly different between high altitude and low altitude groups, multiple regression analysis was used to check association of age and BMI along with altitude and gender for every parameter. However, Age and BMI did not show significant association with any parameter except MCHC (p value > 0.05) after adjusting for the variables mentioned. We observed significant differences (<0.0001) in the RBC, HCT, MCV and Hb concentration between the high and low altitude Tibetans (Figure 2). Mean value of RBC and HCT were higher in high attitude males and females compared low altitude males and females, respectively. Mean value of RBC in high altitude males and low altitude males were 6.21 ± 0.74 and 4.45 ± 0.49 million cells/μl, respectively. In females, these were 5.4 ± 0.40 and 4.19 ± 0.40 cells/μl, respectively. Mean value of HCT in high altitude males, low altitude males, high altitude females and low altitude females were 59.83 ± 6.89, 44.57 ± 4.56, 50.09 ± 3.6, 42.5 ± 4.06 %, respectively. MCV was significantly different only females but not in males with mean value 92.7 ± 6.17 and 101.5 ± 4.67 fL in high altitude and low altitude females respectively. Details of the statistical parameters are given in Table 2 and 3. Significant difference was observed in MCHC also in both males and females, however, age showed significant association with MCHC with coefficient −0.028 (p value 0.00223). Hb concentration was significantly higher in high altitude males and females compared to low altitude males and females. The mean Hb values in males from high and low altitudes were 20.01 ± 1.91 and 13.42 ± 1.75 g/dl, respectively. In females, these were 16.97 ± 1.74 and 12.56 ± 1.19 g/dl, respectively. Details of the tests performed and statistical parameters have been provided in Table 2.

**Figure 2:**
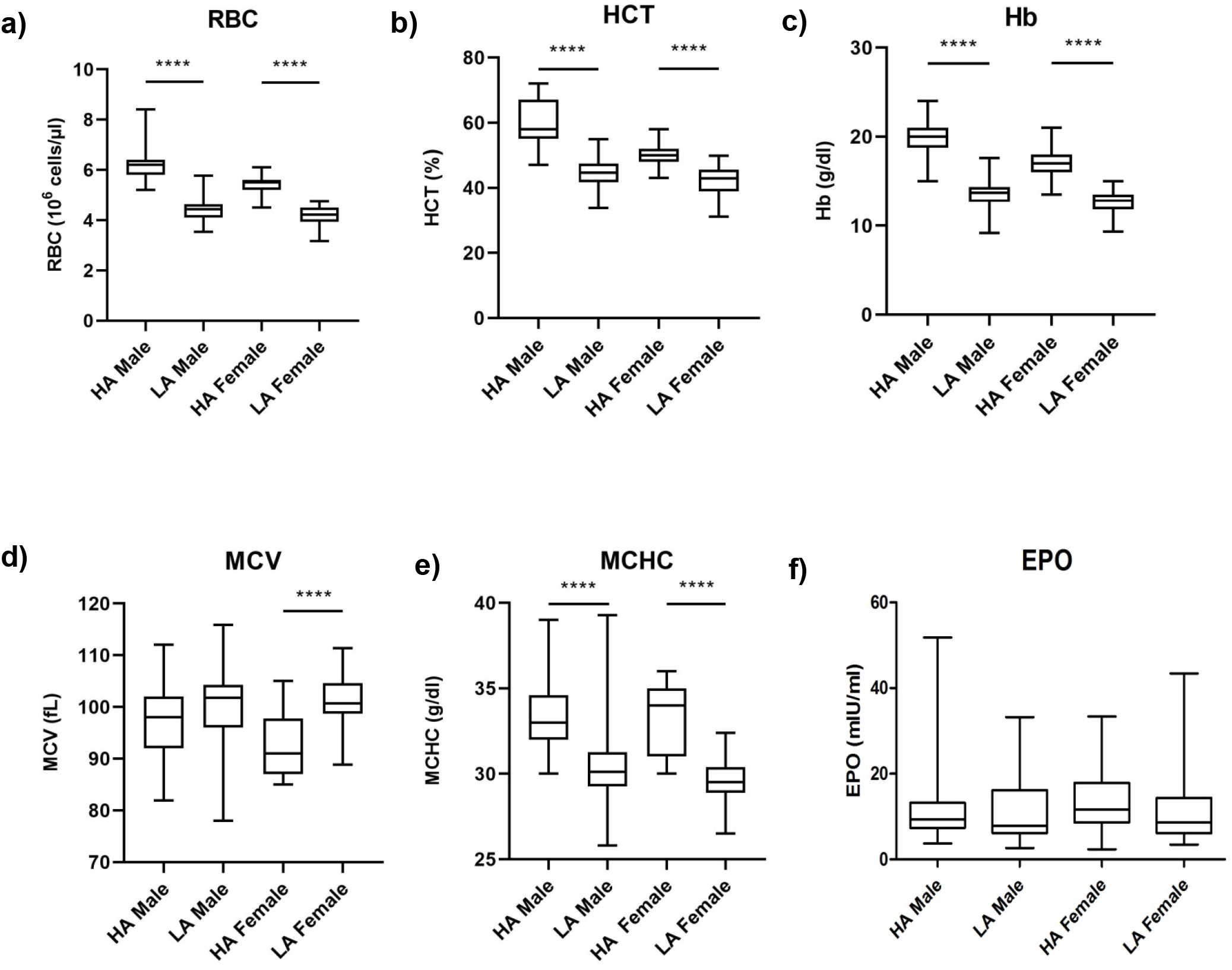
Various hematological parameters of high and low altitude Tibetan males and females: a) RBC b) HCT c) Hb d) MCV e) MCHC f) Serum EPO level

**Table 2:**
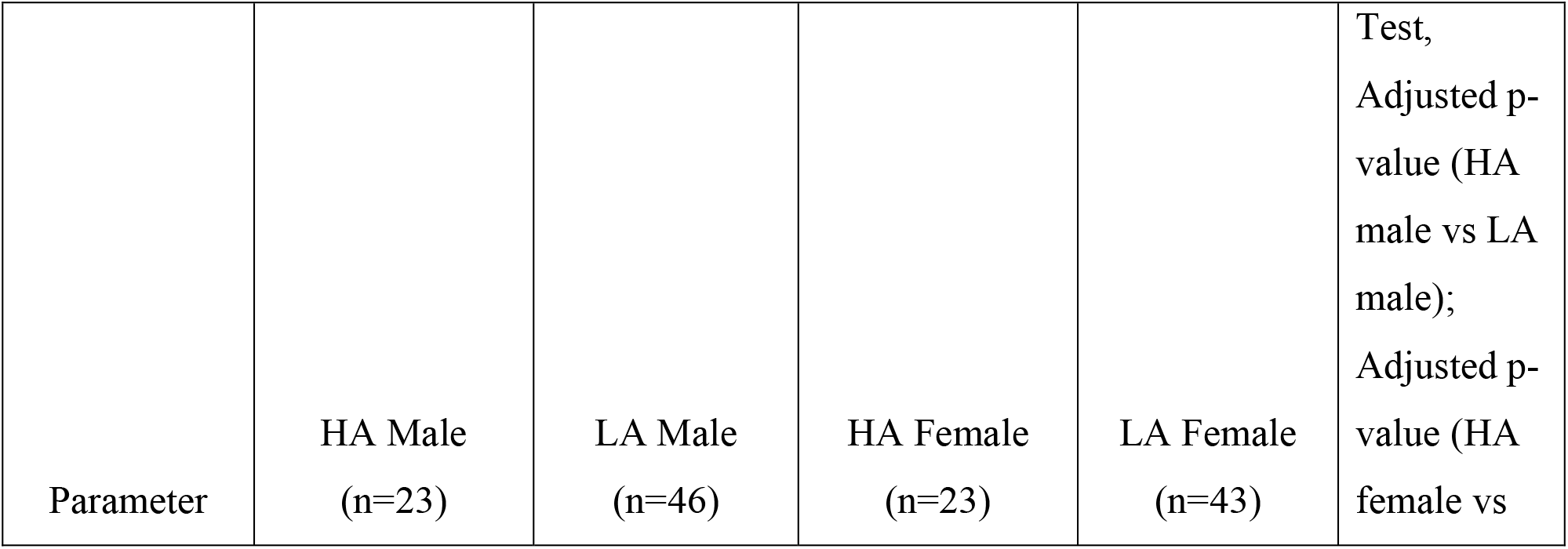
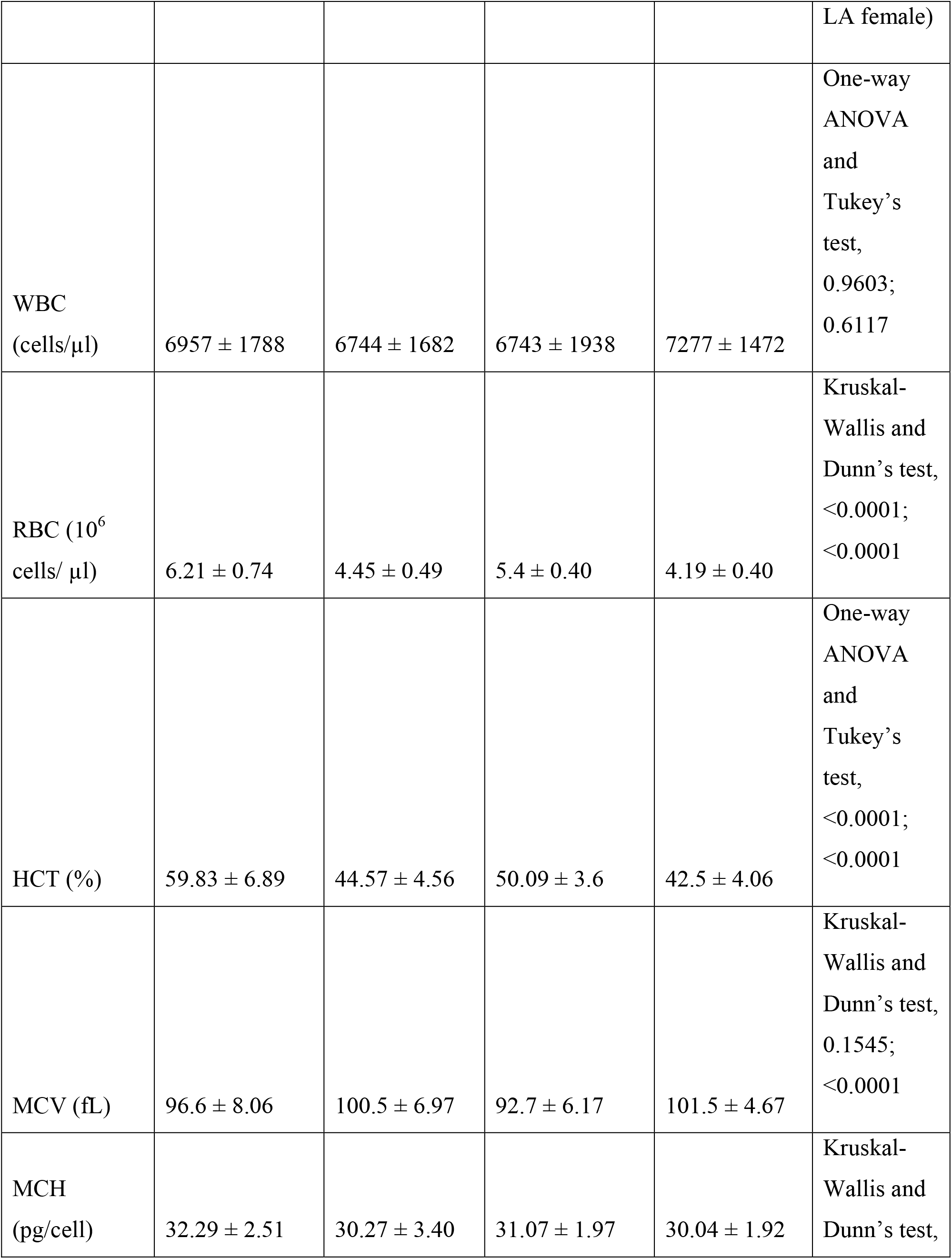
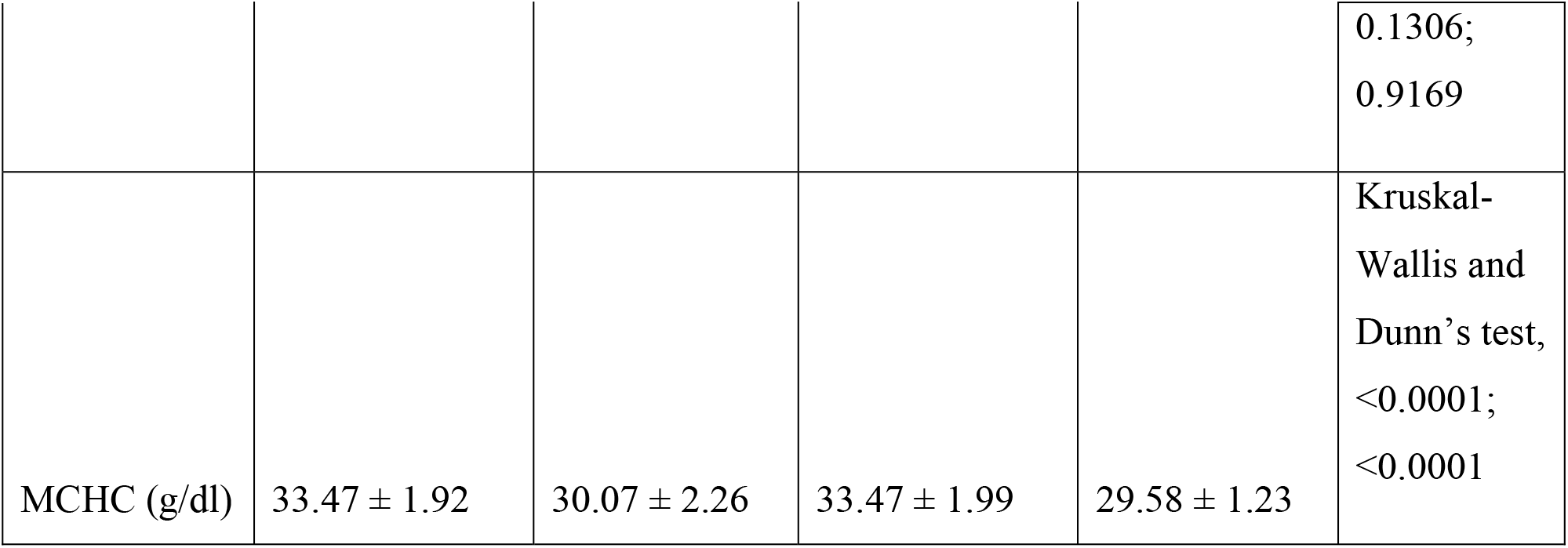
Description of various hematological parameters in the subjects. Data has been expressed in mean ± SD format.

**Table 3:** Description of serum EPO and Hb in the subjects. Data has been expressed in mean ± SD format.

| Parameter | HA Male | LA Male | HA Female | LA Female | Test, Adjusted p-value (HA male vs LA male);<br>Adjusted p-value (HA female vs LA female) |
| --- | --- | --- | --- | --- | --- |
| EPO (mIU/ml) | 13.3 ± 11.61<br>(n=21) | 11.32 ± 7.77<br>(n=42) | 13.21 ± 6.98<br>(n=21) | 11.31 ± 7.63<br>(n=42) | Kruskal-Wallis and Dunn's test,<br>>0.9999; 0.6768 |
| Hb (g/dl) | 20.01 ± 1.91<br>(n=41) | 13.42 ± 1.75<br>(n=46) | 16.97 ± 1.74<br>(n=38) | 12.56 ± 1.19<br>(n=43) | Kruskal-Wallis and Dunn's test,<br><0.0001; <0.0001 |

#### Serum EPO

Serum EPO is one of the prominent outcomes of altitude-induced hypoxia that gets elevated in the people travelling to high altitude. We checked serum EPO level within high and low altitude Tibetan males and females. No significant difference was observed in either males or females with mean value of 13.3 ± 11.61, 11.32 ± 7.77 mIU/ml in males and 13.21 ± 6.98 and 11.31 ± 7.63 mIU/ml in females from high and low altitudes, respectively (Figure 2 and Table 3).

#### Relationship between serum EPO and Hb

We have also assessed the relationship between serum EPO level and Hb (Figure 3a) of the subjects. Both serum EPO and Hb data were available from 130 participants. Interestingly, these parameters did not show any significant correlation (Spearman r = −0.03984, p value = 0.6526).

**Figure 3:**
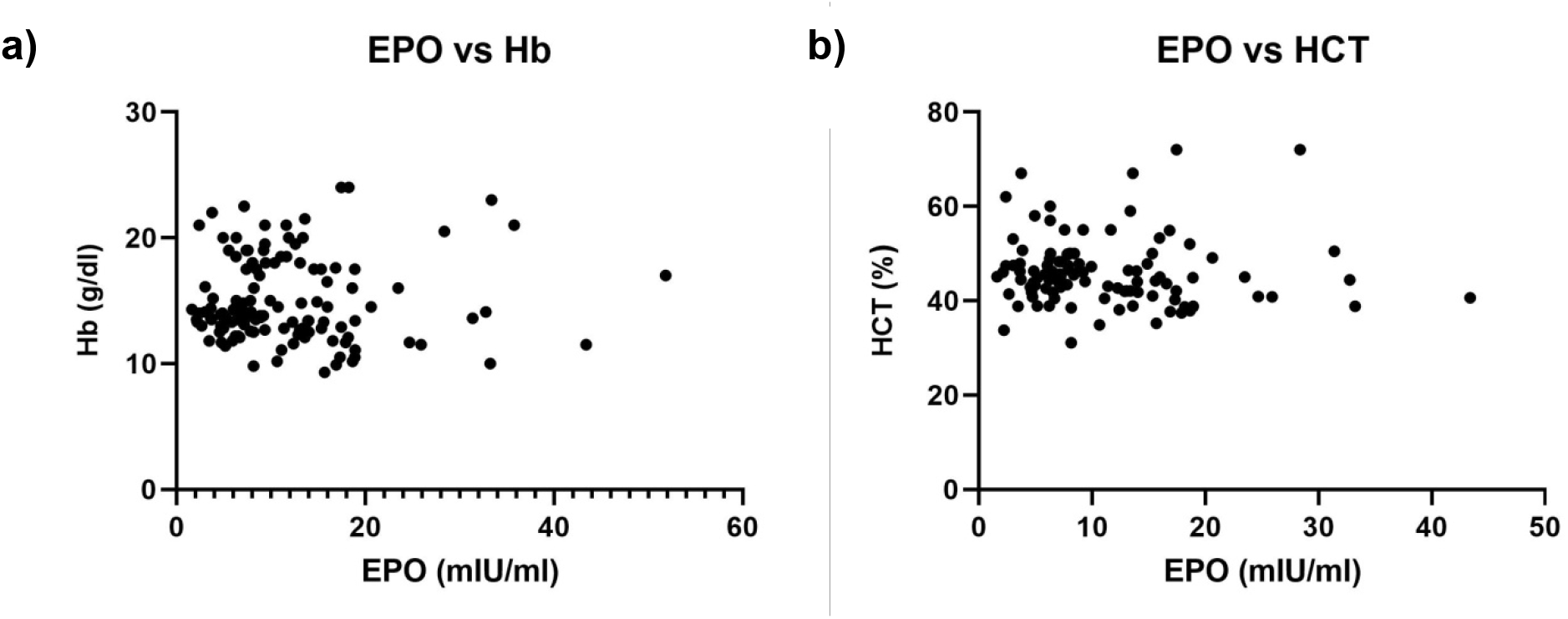
a) Relationship between serum EPO level and Hb and b) serum EPO level and HCT, in Tibetans

Multiple linear regression analysis was performed to check association of Hb with age, gender, altitude and BMI. Among these, only GenderMale and low altitude showed significant effect on hemoglobin concentration. The coefficients were as following-β_0_ (intercept) =17.5462 (p value < 2.25 × 10^-7^), β_1_ (Altitudelow)= −5.4894 (p value < 2.25 × 10^-7^), β_2_ (GenderMale)= 1.9378 (p value = 2.00 × 10^-5^). Adjusted R-squared was 0.7509 (p value < 2.48 × 10^-7^).

#### Relationship between serum EPO and HCT

Next, correlation was assessed between HCT and EPO in the participants, since these two parameters were shown to be correlated in several diseases including renal insufficiency, end-stage renal disease etc ^19,20^. Both serum EPO and HCT data were available from 107 participants. In our study, however, they did not show any significant correlation (Spearman r = 0.1562, p value = 0.1081) (fig.3b).

Multiple linear regression, however, showed association of HCT with gender-male and lowaltitude with coefficients β_0_ (intercept) = 52.5357 (p value < 2.25 × 10^-7^), β_1_ (Altitudelow) = - 11.3502 (p value < 2.25 × 10^-7^), β_2_(GenderMale) = 4.8416 (p value = 1.16 × 10^-3^). Adjusted R-squared was 0.5786 (p value < 2.48 × 10^-7^).

## Discussion

Our study, to the best of our knowledge, shows the difference between Tibetans residing in high and low altitudes for the first time. We observed significantly higher RBC and HCT in high altitude males and females compared to their low altitude counterpart. MCV was significantly higher in low altitude females than high altitude females. Hemoglobin was significantly higher in Tibetans residing in high altitude, compared to low altitude, in both males and females. Our observed range of hemoglobin was little higher than few studies in high altitude Tibetans, however, it was reported that at elevation more than 4500 meters, Hb concentration in Tibetans also show higher mean values ^21–24^. Hb concentration observed in our study for low altitude Tibetans was at par with another study with Tibetans at sea-level ^10^. Serum EPO, in our study, did not show any significant difference in either males or females, that might be due to attenuated erythropoietin response in the population ^25^. A recent study showed that blood cell phenotypes are ancestry dependent and selective pressure can give rise to different blood cell traits ^26^. Our study additionally shows, different exposure of the same population to different environments can also alter blood cell traits. A few studies, however investigated the difference between sea-level Tibetans and other low altitude populations previously, but comparison of low altitude Tibetans with high altitude Tibetans for hematological parameters is not known ^10^. These kinds of studies are important, because, it has been hypothesized by various authors previously, that, a few genetic variants could be very specific to Tibetan population, or, they could moderate hemoglobin concentration, etc., only in high altitude environment ^27^. Our study reveals the difference in various hematological parameters between high and low altitude Tibetans, which goes well with the hypothesis – environment can modulate the expression. Hemoglobin concentration has been shown to play important role in reproductive fitness in Tibetan women, where lower hemoglobin favored reproductive fitness ^28^. It would be interesting to investigate those events separately in the low altitude Tibetans to check, whether the trend is true, since hemoglobin level was lower in the Tibetans from low altitude. Similarly, Tibetan men, having lower hemoglobin concentration are known to show better exercise capacity in high altitude ^21^. Investigation of that between high altitude and low altitude Tibetans would clarify the scenario better, as our study shows; there is striking difference in hemoglobin concentration between both the groups. These kinds of studies would help to understand the physiological aspects better, when native Tibetans are not present in their native high altitude environment as well as to understand the impact of environmental change, knowledge of which could be helpful for tourists and soldiers travelling to high altitude It would be interesting to explore the molecular driving forces behind these kinds of differences. Epigenetic studies, important tool for exploring gene-environment interaction could be fruitful in this context.

## AUTHOR CONTRIBUTIONS

NB, KT designed the study. NB, TN, MMS collected samples. NB, TN performed data acquisition. NB performed wet lab experiments and data analysis. TN, KT provided resources. NB wrote the manuscript with inputs from KT and TN. All the authors critically revised the manuscript and approved the final version of it.

## ACKNOWLEGEMENTS

This work was supported by BSC-0118 (EpiHed) grant provided to KT from the Council of Scientific and Industrial Research (CSIR), Govt. of India (GoI). KT was also supported by J.C. Bose fellowship, Department of Science and Technology (DST), GoI. NB acknowledges DST for DST-INSPIRE fellowship and DST (GoI) and the British Council (UK) for awarding short-term research internship under Newton Bhabha PhD Placement Programme. We thank all the participants of our study. We thank the officers of the Tibetan settlements in Karnataka for their help and co-operating. We thank Nony P Wangchuk, Eashay Lhamo, Ms Sherab Dolma, Tsering Motup, Tsering Palzes, Tsering Dolker, Mrs. Sonam Dolma, Jaison Sequera and Achintya Basak for their help during sample collection and transportation. We thank Dr. Sheikh Nizamuddin for helpful discussions and Dr. Nitin Tupperwar and Dr. Sheikh Nizamuddin for critically reviewing the manuscript.

## Conflict of interest

Authors declare no conflict of interest.

## Notes

### Competing Interest Statement

The authors have declared no competing interest.

